# Communicating mass spectrometry quality information in mzQC with Python, R, and Java

**DOI:** 10.1101/2024.05.06.592686

**Authors:** Chris Bielow, Nils Hoffmann, David Jimenez-Morales, Tim Van Den Bossche, Juan Antonio Vizcaíno, David L. Tabb, Wout Bittremieux, Mathias Walzer

## Abstract

Mass spectrometry is a powerful technique for analyzing molecules in complex biological samples. However, inter- and intra-laboratory variability and bias can affect the data due to various factors, including sample handling and preparation, instrument calibration and performance, and data acquisition and processing. To address this issue, the Quality Control (QC) working group of the Human Proteome Organization’s Proteomics Standards Initiative has established the standard mzQC file format for reporting and exchanging information relating to data quality. mzQC is based on the JavaScript Object Notation (JSON) format and provides a lightweight yet versatile file format that can be easily implemented in software. Here, we present open-source software libraries to process mzQC data in three programming languages: Python, using pymzqc; R, using rmzqc; and Java, using jmzqc. The libraries follow a common data model and provide shared functionality to operate on mzQC files, including the (de)serialization and validation of mzQC files. We demonstrate use of the software libraries for extracting, analyzing, and visualizing QC metrics from different sources and show how these libraries can be integrated with each other, with existing software tools, and in automated workflows for the QC of mass spectrometry data. All software libraries are available as open source under the MS-Quality-Hub organization on GitHub (https://github.com/MS-Quality-Hub).

## Introduction

Mass spectrometry (MS) is a powerful analytical technique for analyzing molecules in complex biological samples. However, MS data are inherently prone to variability and bias. Even between related MS experiments, subtle variations in sample preparation, instrument performance, and data processing can lead to hidden data inconsistency.^1^ To inspire confidence in the results of an MS experiment and to ensure that different measurements are consistent and comparable, it is of key importance that appropriate quality assurance and quality control (QC) strategies are taken to monitor and control the existing variability. Therefore, QC is essential for generating high-quality MS data that can support meaningful and reproducible scientific discoveries. This is especially relevant in light of the reproducibility crisis in science.^2^ By improving the quality assessment and reproducibility of MS experiments, QC ensures the credibility and confidence of the resulting scientific findings.

Unfortunately, so far no unified approaches towards QC in biological MS have been established. One of the limitations is the lack of a standard file format to store and communicate QC metrics, which are numerical or graphical indicators that describe the quality of MS data at different levels, such as sample quality, instrument performance, completeness of the measurements, and data consistency.^3^ Currently, QC metrics are often stored in different formats and locations, such as instrument log files, proprietary software outputs, and spreadsheets. This makes it difficult to access, compare, and share QC information over time, across different instruments, sample preparation techniques, and laboratories.

To address this issue, the Quality Control working group^4^ of the Human Proteome Organization’s Proteomics Standards Initiative (HUPO-PSI)^5^ has recently established the standard mzQC file format (https://github.com/HUPO-PSI/mzQC) to report and exchange data quality-related information for MS experiments and the associated analysis results. mzQC is based on the widespread JavaScript Object Notation (JSON) format to provide a lightweight yet versatile file format that can be easily implemented in software to produce or consume mzQC files, and its goal is to support diverse workflows in proteomics, metabolomics, and other MS applications. It is important to note that mzQC aims to provide a standardized framework for storing and exchanging QC metrics in MS data analysis, rather than to directly judge the quality of the data it describes.

QC metrics in an mzQC file are grouped in “`runQuality`” or “`setQuality`” elements, depending on whether the metrics pertain to a single or multiple MS runs, respectively. Each `runQuality` or `setQuality` element contains a “`metadata`” section that provides information to track the provenance of the QC metrics, such as the originating MS run(s) and the software tool(s) used to calculate the metrics. QC metric values are stored in “`qualityMetric`” elements and can consist of single values, tuples, or tabular data. Additionally, each QC metric is defined by a corresponding term in the PSI-MS controlled vocabulary^6^ for semantic annotation of the data and to ensure an unambiguous definition of each QC metric. Further technical details of the mzQC format and the official PSI specification document (version 1.0, released in February 2024) are available at https://github.com/HUPO-PSI/mzQC.

To ensure the adoption of the mzQC format, supporting software tools are needed. There is a vibrant open-source community of bioinformaticians developing software to analyze MS data in various programming languages, among which some of the most popular are Python,^7–10^ which is widely used for data analysis and machine learning; R,^11,12^ a language designed for statistical computing and graphics; and Java,^13,14^ a multi-platform programming language that is suitable for large-scale applications.

In this manuscript, we present open-source software libraries to read, write, and validate QC data in the mzQC format in the three programming languages mentioned above: Python, R, and Java. We describe the design and implementation of these libraries, which follow a common data model and provide shared functionality to operate on mzQC files. We demonstrate the use of these software libraries for extracting, analyzing, and visualizing QC metrics from different sources. We also show how these libraries can be integrated with existing software tools and workflows for performing QC of MS data, with mzQC acting as the glue between various workflow steps. All software libraries are available as open source under the MS-Quality-Hub organization on GitHub (https://github.com/MS-Quality-Hub/).

## Methods

### mzQC software libraries

The mzQC software libraries are implemented in three popular programming languages (Table 1): pymzqc in Python, rmzqc in R, and jmzqc in Java. Each library builds on the mzQC schema definition, which formally defines the syntax of mzQC documents using a JSON schema, and provides a high-level abstraction of data quality-related information in mzQC files. Rather than a single application programming interface (API) that all libraries share, they each follow the conventions and best practices of their respective programming languages.

**Table 1.**
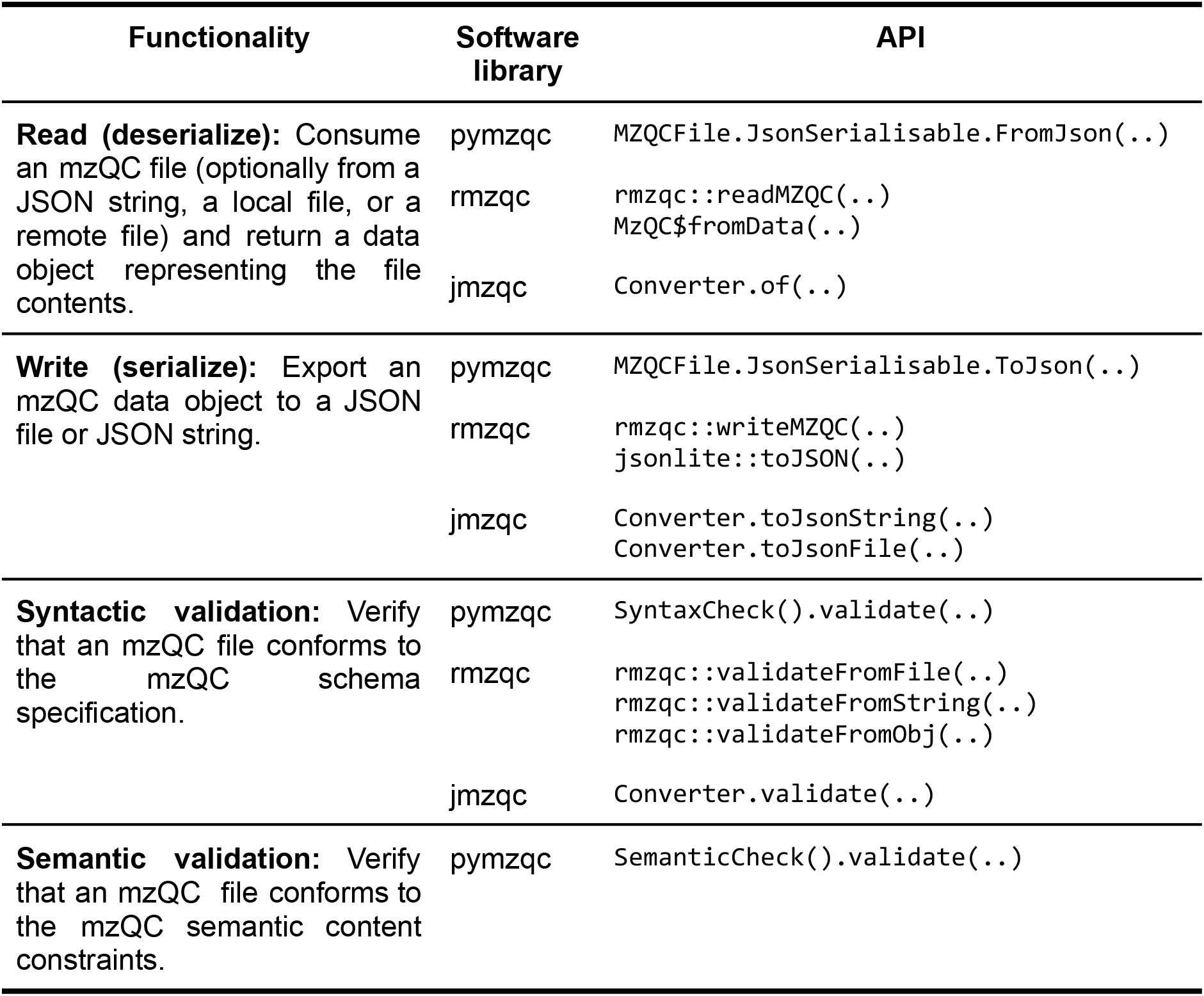
Overview of the high-level functionality provided by the mzQC software libraries.

The primary functionality provided by all three software libraries is the serialization and deserialization of mzQC files, which facilitates the reading and writing of QC information, respectively. This enables users to create mzQC files containing newly computed QC information, read existing mzQC files with previously computed QC metrics, and manipulate the QC information for further data processing and analysis. The software libraries automatically perform native value type matching where possible, such as converting tabular data to data.frame objects in R or Pandas DataFrame objects in Python.

In addition to (de)serialization, the software libraries provide user-friendly functionality to validate mzQC files. Syntactic validation checks if the structure of mzQC data conforms to the defined syntax rules, ensuring that the data are structured correctly and contain all necessary pieces of information. Semantic validation, on the other hand, involves verifying that the data make sense in their specific context, ensuring that they meaningfully and logically represent MS-related QC concepts and information. All three libraries support syntactic validation, which is based on the mzQC JSON schema. The pymzqc library also supports semantic validation, which interprets the content of mzQC files to ensure the correctness of the QC information, including verification that all QC metrics are represented in an accessible controlled vocabulary or ontology and that the data value types match the definition in the controlled vocabularies. Additionally, a web application to validate mzQC files, powered by pymzqc, is available at https://hupo-psi.github.io/mzQC/validator/.

### Code availability

All mzQC supporting software libraries are freely available on GitHub as open source (Table 2), collected in the MS-Quality-Hub organization (https://github.com/MS-Quality-Hub). Additionally, all software libraries can be easily installed using their respective language-preferred toolchains (Table 2). All software libraries follow development best practices, including extensive code documentation, detailed installation instructions, and automated testing using continuous integration.

**Table 2.**
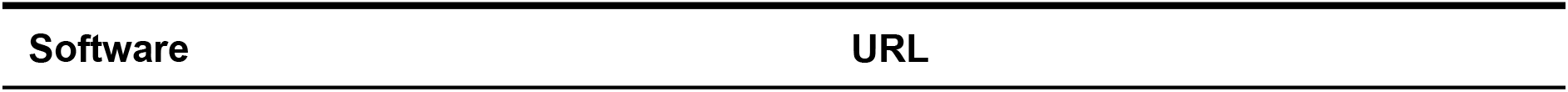
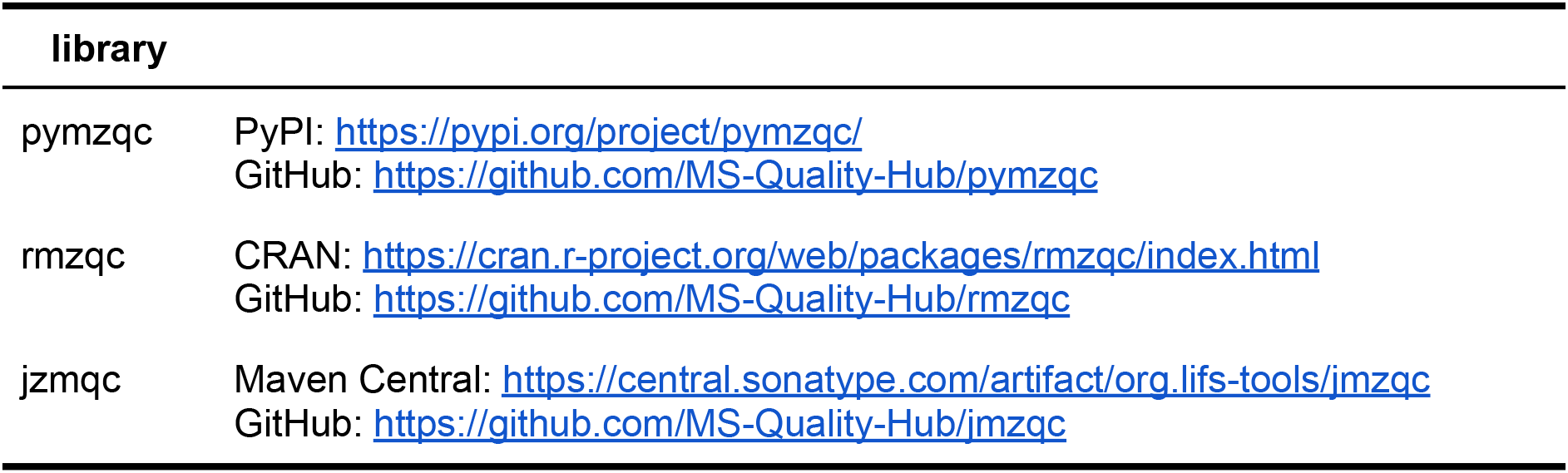
Availability of the software libraries in their respective software package and source code repositories.

| Software | URL |
| --- | --- |

| library |  |
| --- | --- |
| pymzqc | PyPI: https://pypi.org/project/pymzqc/<br>GitHub: https://github.com/MS-Quality-Hub/pymzqc |
| rmzqc | CRAN: https://cran.r-project.org/web/packages/rmzqc/index.html<br>GitHub: https://github.com/MS-Quality-Hub/rmzqc |
| jzmqc | Maven Central: https://central.sonatype.com/artifact/org.lifs-tools/jmzqc<br>GitHub: https://github.com/MS-Quality-Hub/jmzqc |

All code used in this manuscript to demonstrate the library’s capabilities can be found as open source in a dedicated GitHub repository under the MS-Quality-Hub organization at https://github.com/MS-Quality-hub/mzqclib-manuscript.

### Data

We have reanalyzed MS data from a proteomics study of anaerobic respiration in *E. coli* grown in sulforaphane, obtained via ProteomeXchange^15^ with dataset identifier PXD040621.^16^ In this study, bacterial cultures were grown in the presence of either sulforaphane (10 μM) or 0.034% dimethyl sulfoxide (DMSO) as a control. The study comprised four biological replicates of bacterial growth in both conditions, acquired using a 120 min liquid chromatography gradient measured on an Orbitrap Q-Exactive using data-dependent acquisition.

The data were reanalyzed by converting the raw files to mzML^17^ using ThermoRawFileParser (biocontainer version 1.4.0)^18^ and sequence database searching using Tide (Crux toolkit version 4.2).^19^ We performed target–decoy searching against the UniProtKB^20^ *E. coli K12* reference proteome (UP000000625, downloaded on October 20, 2023), and configured the search for tryptic peptides with up to three missed cleavages, variable oxidation of methionine, and a precursor mass tolerance of 50 ppm. Spectrum identifications were filtered at a 1% protein-level false discovery rate with crema (version 0.0.10).^21^

## Results

To illustrate the functionality and interoperability of the mzQC software libraries, we first used each library separately to produce mzQC files containing various QC metrics, after which the individual mzQC files were combined into a single QC report. This demonstration deliberately splits QC metric calculation and mzQC file generation across three programming languages to demonstrate the interoperability of the software libraries.

Our example workflow consists of several steps (Figure 1). First, the raw files were converted to mzML and processed using sequence database searching. Next, various QC metrics (Supplementary Table 1) were computed using dedicated scripts in Java, Python, and R and exported to mzQC files using the mzQC software library for the corresponding programming language:

**Figure 1:**
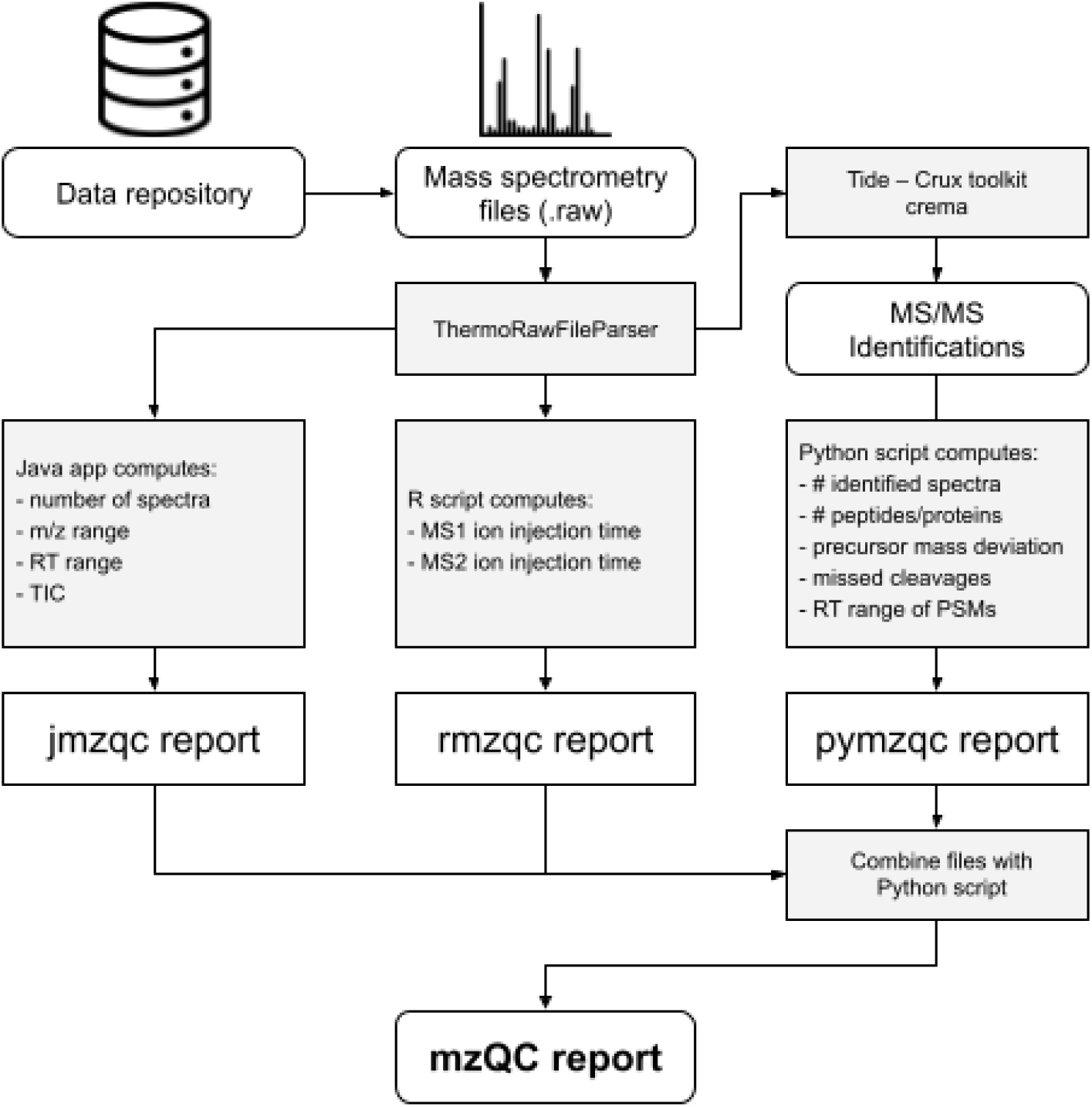
mzQC processing workflow. Each software library is separately used to process different QC metrics, which are ultimately combined into a single QC report.

### jmzqc

As a compiled language, Java is excellently suited to process large amounts of data. Consequently, we used jmzqc combined with jmzml^22^ and MSDK^23^ to efficiently read the mzML peak files and compute basic QC metrics from the MS data. We calculated the number of chromatograms, the *m*/*z* range of the acquired spectra, the retention time range of the acquired spectra, the total ion chromatogram, and the base peak intensities.

### rmzqc

Based on R’s emphasis on statistical processing, we used rmzqc to collect statistics of the ion injection parameters at the level of MS and MS/MS spectra.

### pymzqc

We used pymzqc in combination with Pyteomics^8^ and Pandas^24^ to read the peak and identification data; count the number of MS/MS spectra, the number of identified MS/MS spectra, the number of identified peptides, and the number of identified proteins; compute the distribution of the precursor mass deviation of the identifications; evaluate the number of missed cleavages; and find the retention time range during which spectra could be successfully annotated.

Finally, pymzqc was used to combine the mzQC outputs obtained during each step and merge the data into a single mzQC file. A Jupyter notebook^25^ was used for subsequent interactive data analysis and to produce a final report that summarizes the QC metrics. Metrics across all MS runs were visualized using a clustered heatmap, with metric values percentile rank scaled (Figure 2).

**Figure 2:**
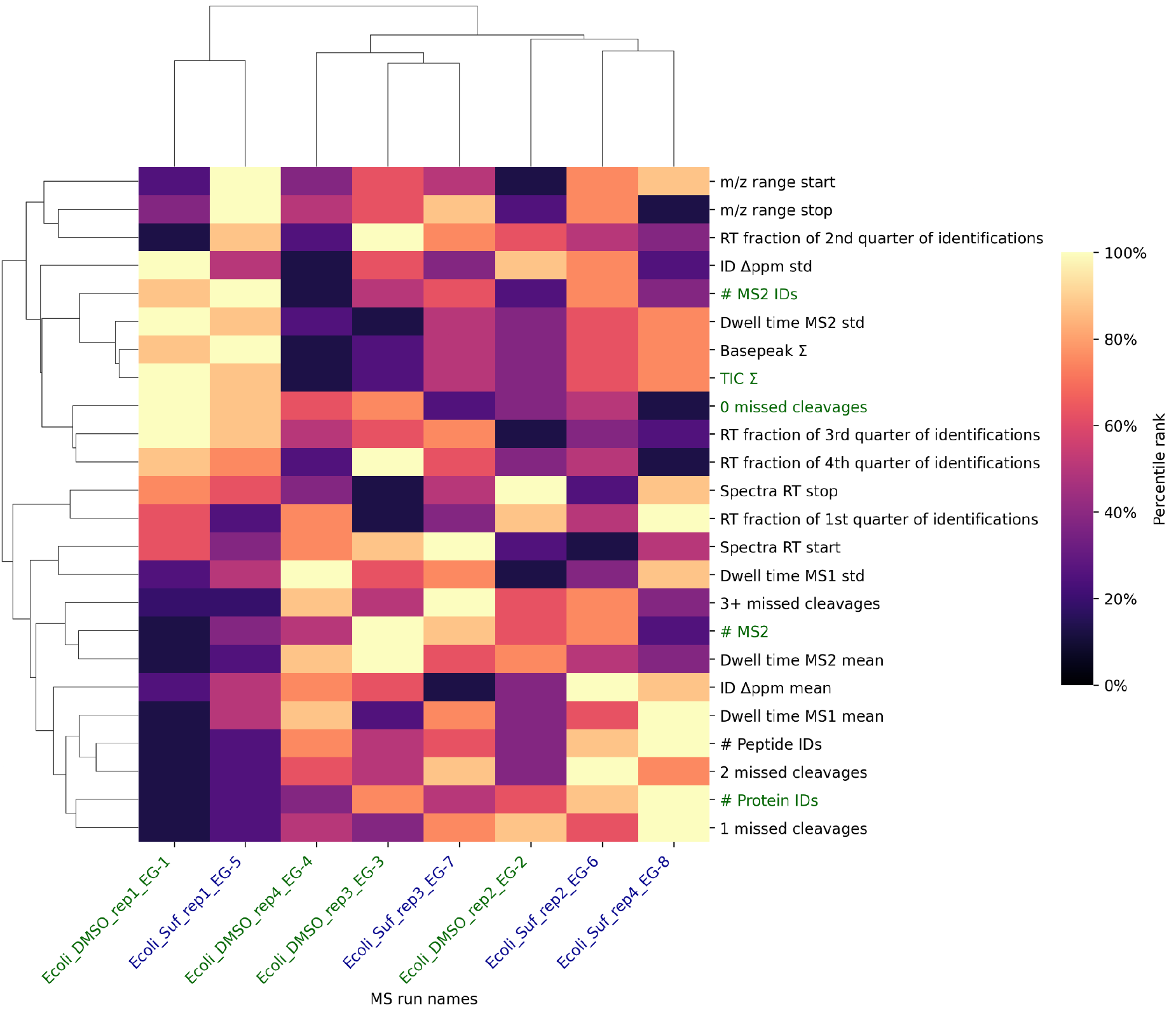
Heatmap with dendrogram displaying QC metrics across eight MS runs (DMSO controls colored in green, sulforaphane samples colored in blue), clustered by MS runs on the horizontal axis and by QC metrics on the vertical axis. Colors in the heatmap represent percentile ranks calculated from the combined dataset, with darker shades indicating lower percentile ranks and lighter shades indicating higher ranks. The QC metrics include the number of acquired MS/MS spectra, MS/MS identifications, peptide identifications, protein identifications, summed total ion current, and number of missed cleavages, among others. The metrics discussed in the text are highlighted in green. See the Jupyter Notebook on our GitHub repository (https://github.com/MS-Quality-Hub/mzqclib-manuscript/) for data analysis and code to generate the plot.

As a brief example, we performed a visual inspection of the QC metrics to illustrate how QC data can be used to explore the implications of MS data quality (Figure 2). Note that this description is not provided by mzQC directly, but is based on the author’s interpretation of the heatmap. When examining the heatmap and its clustering of QC metrics across the eight MS runs, we observe a distinct separation between two specific runs from the rest: the Ecoli_DMSO_rep1_EG-1 control run, and the Ecoli_Suf_rep1_EG-5 sulforaphane run. This divergence is driven by a comparatively lower number of MS/MS spectra acquired, peptides and proteins identified, as well as related QC metrics. Interestingly, these two runs exhibit above average total ion currents and the rate of MS/MS spectra that could be identified, suggesting that the lower peptide and protein identification rate is due to a decrease in the number of MS/MS spectra acquired, rather than due to a reduction in the quality of the MS/MS spectra. Additionally, the two outlier runs have a higher proportion of peptides with no missed tryptic cleavages, which might impact the subsequent protein inference. Consequently, deriving biological interpretations from the full experiment may require robust statistical models that are resistant to outliers.

The large contrast in the number of acquired MS/MS spectra and identifications indicates that some caution might need to be exercised when interpreting the results for these two runs to study the effect of sulforaphane on *E. coli* growth. The heatmap generated by the mzQC pipeline suggests the need for a deeper investigation into the cause of these discrepancies. This will necessitate a broader QC analysis incorporating long-term instrument performance monitoring, including repeated analysis of consistent QC samples. This would provide a more informed basis for interpreting these outlier results within the context of the study.

## Conclusion

The introduction of the mzQC standard file format for quality control in biological mass spectrometry has numerous potential benefits, including increased reproducibility, improved interoperability, and enhanced data sharing among researchers. However, adopting new file formats can be challenging without suitable software libraries to facilitate their integration into bioinformatics software. The development of open software libraries such as pymzqc, rmzqc, and jmzqc is therefore essential for the successful adoption of mzQC. These libraries provide a consistent interface for accessing mzQC files and allow bioinformatics software developers to easily incorporate mzQC into their tools and workflows. Third-party support for mzQC is already emerging,^26^ and will be bolstered further by the presented software libraries.

Based on a worked use case, we have demonstrated that only a small amount of code is needed to construct an mzQC object in memory, populate it with calculated metric values, and export the data to an mzQC file on disk. Likewise, the interactive notebooks showcase the libraries for conveniently reading data from mzQC for further processing and reporting. This illustrates how the high-level abstractions provided by the mzQC libraries presented here facilitate interacting with mzQC files in different programming languages.

The mzQC software libraries offer several key benefits, including the ability to validate mzQC files, extract information from them, and convert to and from other formats. Additionally, the complexity of the mzQC software libraries is limited, building on native JSON support in the various programming languages. These capabilities are crucial for the development of new QC tools and workflows that can help to improve the reliability and reproducibility of mass spectrometry experiments. Especially as data analysis pipelines are increasingly gaining in complexity, with separate processing steps implemented in different programming languages instead of a single monolithic application, this multi-language support will be highly beneficial. As such, we anticipate that our software libraries will foster a vibrant ecosystem of general and bespoke bioinformatics tools for QC of MS experiments, which will be able to seamlessly interoperate through a common mzQC interface.

In light of these benefits, we invite software developers to start using the mzQC software libraries. Additionally, we welcome any contributions to the mzQC software libraries and the mzQC format, for example by developing complementary libraries in alternative programming languages, such as C++, C#, Rust, or JavaScript. By doing so, the adoption of mzQC in the mass spectrometry and bioinformatics communities will be further facilitated, ultimately leading to better-quality data and more reliable scientific discoveries. To ensure the continued evolution and application of mzQC, we are dedicated to enhancing its ecosystem, including broadening its integration into diverse bioinformatics tools and developing extensive use cases for QC across various biological mass spectrometry applications.

## Acknowledgements

M.W. would like to acknowledge funding from the H2020 EPIC-XS grant [grant number 823839], BBSRC ‘Proteomics DIA’ [BB/P024599/1], and from The Wellcome Trust [208391/Z/17/Z]. Additionally, J.A.V. would like to acknowledge EMBL core funding. This work was supported in part by the de.NBI Cloud within the German Network for Bioinformatics Infrastructure (de.NBI) and ELIXIR-DE (Forschungszentrum Jülich and W-de.NBI-001, W-de.NBI-004, W-de.NBI-008, W-de.NBI-010, W-de.NBI-013, W-de.NBI-014, W-de.NBI-016, W-de.NBI-022). T.V.D.B. acknowledges funding from the Research Foundation Flanders (FWO) [1286824N]. W.B. acknowledges support by the University of Antwerp Research Fund.

## Supplementary Information

**Supplementary Table 1:** QC metrics considered during the analysis and their accession numbers in the PSI-MS controlled vocabulary. The last column refers to the programming language used to compute the metrics.

| Accession | Metric | Programming Language |
| --- | --- | --- |
| MS:1000505 | base peak intensity | Java |
| MS:1002404 | count of identified proteins | Python |
| MS:1003044 | number of missed cleavages | Python |
| MS:1003250 | count of identified peptidoforms | Python |
| MS:1003251 | count of identified spectra | Python |
| MS:4000060 | number of MS2 spectra | Python |
| MS:4000069 | m/z acquisition range | Java |
| MS:4000070 | retention time acquisition range | Java |
| MS:4000071 | number of chromatograms | Java |
| MS:4000104 | total ion currents | Java |
| MS:4000132 | MS1 ion collection time mean | R |
| MS:4000133 | MS1 ion collection time sigma | R |
| MS:4000137 | MS2 ion collection time mean | R |
| MS:4000138 | MS2 ion collection time sigma | R |
| MS:4000178 | precursor ppm deviation mean | Python |
| MS:4000179 | precursor ppm deviation sigma | Python |
| MS:4000180 | table of missed cleavage counts | Python |
| MS:4000181 | identified MS2 quarter RT fraction | Python |

## Notes

### Competing Interest Statement

The authors have declared no competing interest.

